# Porcine pancreatic ductal epithelial cells transformed with KRAS^G12D^ and SV40T are tumorigenic

**DOI:** 10.1101/2021.01.15.425792

**Authors:** Katie Bailey, Sara B. Cartwright, Neesha S. Patel, Neeley Remmers, Audrey J. Lazenby, Michael A. Hollingsworth, Mark A. Carlson

**Affiliations:** Department of Surgery, University of Nebraska Medical Center, Omaha NE 68198, USA; VA Medical Center, Omaha NE 68105, USA; Department of Pathology and Microbiology, University of Nebraska Medical Center, Omaha NE 68198, USA; Eppley Institute for Research in Cancer, University of Nebraska Medical Center, Omaha NE 68198, USA; Fred & Pamela Buffett Cancer Center, University of Nebraska Medical Center, Omaha NE 68198, USA; Department of Genetics, Cell Biology and Anatomy, University of Nebraska Medical Center, Omaha NE 68198, USA; Center for Advanced Surgical Technology, University of Nebraska Medical Center, Omaha NE 68198, USA

**Author notes:** Corresponding author: Mark A. Carlson, Surgery 112, VAMC, 4101 Woolworth Ave Omaha, NE, USA.

## Abstract

We describe our initial studies in the development of an orthotopic, genetically-defined, large animal model of pancreatic cancer. Primary pancreatic epithelial cells were isolated from pancreatic duct of domestic pigs. A transformed cell line was generated from these primary cells with oncogenic KRAS and SV40T. The transformed cell lines outperformed the primary and SV40T immortalized cells in terms of proliferation, population doubling time, soft agar growth, transwell migration and invasion. The transformed cell line grew tumors when injected subcutaneously in nude mice, forming glandular structures and staining for epithelial markers. Future work will include implantation studies of these tumorigenic porcine pancreatic cell lines into the pancreas of allogeneic and autologous pigs. The resultant large animal model of pancreatic cancer could be utilized for preclinical research on diagnostic, interventional, and therapeutic technologies.

## Introduction

In the United States in 2020, approximately 57,600 people will be diagnosed with pancreatic cancer (~3.2% of all new cancer diagnoses), and there will be ~47,000 deaths from pancreatic cancer (~7.8% of all cancer deaths)^1^. The lifetime risk for pancreatic cancer is approximately 1 in 64^2^. The incidence of pancreatic cancer has been gradually increasing since the mid-1990’s, and generally is higher in the African-American population^3^. Pancreatic cancer is now the fourth most common cause of cancer-related death in both men and women (after lung, prostate, and colorectal cancer, or lung, breast, and colorectal cancer, respectively)^1^. Despite advances in treatment modalities and strategies^4^, the mortality rate from pancreatic cancer has not decreased^1^. The overall 5-year survival rate from pancreatic cancer in the U.S. is 9%; 5-year survival rates in localized, regional (nodal spread), or metastatic disease were 37, 12, and 3%, respectively^1^. So, there remains a need for improved early diagnosis and therapy for pancreatic cancer.

Rodent models of pancreatic cancer may not accurately reflect human biology because of differences in anatomy, physiology, immune response, and genetic sequence between the two species^5–9^. Remarkably, only 5-8% of anti-cancer drugs that emerged from preclinical studies and entered clinical studies were ultimately approved for clinical use^10,11^. The cause of this low approval rate is multifactorial, but likely includes the less-than-optimal predictive ability of some murine models (e.g., tumor xenografting into immunosuppressed mice) to determine the efficacy of various therapeutics in humans^5–7,12–16^. Moreover, there are a number of genes for which the genotype-phenotype relationship is discordant between mice and human, including *CFTR*^−/−^ and *APC*^+/−17,18^. Incidentally, both the porcine *CFTR*^−/−^ and *APC*^+/−^ mutants reiterate the human phenotype (pulmonary/GI disease and rectal polyposis, respectively)^17–19^, in contradistinction to the murine mutants.

The recent trend to employ genetically-engineered mouse models (GEMM), patient-derived xenografts (PDX), humanized mice, and *in vivo* site-directed CRISPR/Cas9 gene-edited mice in the testing of anti-cancer therapeutics may yield murine models with better predictive ability than obtained previously^7,20–24^. Though promising, these more advanced murine models come with increased cost and complexity^22^, and experience with them still is early. Importantly, all murine models have limited utility in the development of diagnostic or interventional technology that requires an animal subject whose size approximates a human. So, at present, there remains a need for improved animal models of pancreatic cancer that (1) are more predictive of human response to anti-cancer therapy^22,24^, and (2) are of adequate size for development of some diagnostic or interventional technologies. Herein we describe some initial steps taken to develop an orthotopic porcine model of PDAC through genetic manipulation of key genes associated with PDAC within cultured porcine pancreatic ductal epithelial cells.

## Materials and Methods

### Standards, rigor, reproducibility, and transparency

When feasible, the animal studies of this report were designed, performed, and reported in accordance with both the ARRIVE Guidelines (Animal Research: Reporting of *In Vivo* Experiments^25^) and the National Institutes of Health Principles and Guidelines for Reporting Preclinical Research^26,27^; for details, refer to **Fig. S1** and **Table S1**, respectively.

### Materials, reagents, and animal subjects

All reagents were purchased through Thermo Fisher Scientific (www.thermofisher.com) unless otherwise noted. Short DNA sequences for vector construction, mutagenesis, and amplification purposes are shown in **Table S2**. Antibody information is given in **Table S3**. Athymic homozygous nude mice (Nu/J, Foxn1^nu^; females; 4 weeks old) were purchased from The Jackson Laboratory (www.jax.com). DNA sequencing was performed by the UNMC Genomics Core Facility (www.unmc.edu/vcr/cores/vcr-cores/genomics).

### Animal welfare

The animals utilized in this report were maintained and treated in accordance with the *Guide for the Care and Use of Laboratory Animals* (8^th^ ed.) from the National Research Council and the National Institutes of Health^28^, and also in accordance with the Animal Welfare Act of the United States (U.S. Code 7, Sections 2131 – 2159). The animal protocols pertaining to this manuscript were approved by the Institutional Animal Care and Use Committee (IACUC) of the University of Nebraska Medical Center (ID number 16-133-11-FC). All procedures were performed in animal facilities approved by the Association for Assessment and Accreditation of Laboratory Animal Care International (AAALAC; www.aaalac.org) and by the Office of Laboratory Animal Welfare of the Public Health Service (grants.nih.gov/grants/olaw/olaw.htm). All surgical procedures were performed under isoflurane anesthesia, and all efforts were made to minimize suffering. Euthanasia was performed in accordance with the 2013 AVMA Guidelines for the Euthanasia of Animals^29^.

### Isolation of porcine pancreatic ductal epithelial cells

A detailed protocol for isolation of porcine pancreatic ductal epithelial cells (PDECs) is provided in **Fig. S2**. In brief, the intact pancreas from male domestic swine (age 5 mo) was harvested within 5 min after euthanasia, which was accomplished by transection of the intrathoracic inferior vena cava and exsanguination while under deep isoflurane anesthesia. The donor pigs had been on a separate research protocol (a study biomaterials within dorsal skin wounds), and had not received any recent medication other the anesthetics given for euthanasia; buprenorphine and cefovecin sodium had been given 4 weeks prior to euthanasia. Immediately after explantation of the pancreas, the main pancreatic duct was dissected sterilely with microsurgical instruments from the organ body, using 3.5x loupe magnification. A 3-4 cm segment of proximal duct, from the duodenal ampulla to the mid-portion of the duodenal lobe of the porcine pancreas, was isolated and cleared of loose connective tissue. The duct was then minced with microscissors, and enzymatically digested in 10 mL of Collagenase I (10 mg/mL) plus DNase I (100 ug/mL) for 30 min at 37°C, including mechanical disruption through a pipette tip after every 10 min of incubation. After 30 min the cells were pelleted (500 *g* × 5 min), the supernatant was discarded, and the cell pellet was resuspended in Epi Cell Growth Medium (ECM; Sigma, cat. no. 215-500) and pelleted a second time. The cells were then resuspended in 2 mL of complete medium, which was defined as ECM with 5% FBS and 1% Antibiotic-Antimycotic Solution (Corning Inc., cat. no. 30-004-CI), and incubated in a 6 well plates that were precoated with porcine gelatin (Corning Inc., cat. no. 354652). After 4 hours the medium was transferred to a new well and 2 mL of fresh complete medium was added to the previous well. Epithelial colonies started to form after 5-7 days of culture under standard conditions (complete medium, 37°C, 5% CO_2_). If fibroblasts were present on inverted phase microscopy, then light trypsinization was used to remove the contaminating fibroblasts. Light trypsinization consisted of a 3 min incubation with 1 mL of trypsin (Trypsin-EDTA 0.25%, Gibco™/ ThermoFisher, cat. no. 25200056), followed by washing with complete medium, and finishing with addition of 2 mL of the same. Standard trypsinization (detaching all cells for passaging) required a 10-15 min incubation. Cells were passaged in this fashion at least four times using 6-well plates, until no cells with fibroblast morphology were present. The resulting epithelial cells were passaged one time in 10 cm dishes prior to the below use.

### PDEC immortalization & KRAS vector

Primary PDECs were immortalized with SV40 large T antigen, using ready-to-use Lenti-SV40T (Puro) Lentivirus, High Titer cell immortalization kit (Applied Biological Materials, cat. no. LV613; www.abmgood.com), per the manufacturer’s instructions. The source of the porcine *KRAS*^G12D^ mutant was the plasmid used to generate the p53/KRAS Oncopig^30,31^. The *KRAS*^G12D^ cDNA was amplified out of this plasmid with primers (see **Table S2**) that flanked the sequence with *Xho*I and *Bam*HI restriction sites. The amplified product was inserted into the TOPO^®^ vector (TOPO^®^ TA Cloning^®^ Kit; Invitrogen™/ Life Technologies™, Thermo Fisher Scientific, cat. no. K202020) and verified by sequencing, as described above. The pLVX-IRES-ZsGreen1 (Takara Bio/ Clontech Laboratories, cat. no. 632187; www.takarabio.com) vector was cut with *Xho*I and *Bam*HI, and the *KRAS*^G12D^ sequence was ligated into this plasmid, producing a pLVX-IRES-ZsGreen1 vector which contained the mutant cDNA within its multiple cloning site (*KRAS*^G12D^ upstream).

The newly-constructed plasmid, hereafter designated as LV-GK (LV-lentiviral; G = ZsGreen1; K = KRAS^G12D^), was transformed into One Shot™ Stbl3™ Chemically Competent *E. coli* (Invitrogen™/Thermo Fisher Scientific, cat. no. C737303), and plasmid DNA subsequently was isolated using a QIAGEN Plasmid Maxi Kit (cat. no. 12162), all per the manufacturer’s instructions. This plasmid then was transfected into Lenti-X™ 293T cells, using Takara’s Lenti-X (VSV-G) Packing Single Shots (Takara Bio/ Clontech Laboratories, cat. no. 631275, www.takarabio.com; per the instructions), to generate infectious lentiviral particles that would produce direct expression of ZsGreen1 and KRAS^G12D^ mutant in the transduced cells. The medium for the Lenti-X™ 293T cells was DMEM + 10% tetracycline-free FBS (Gemini Bio-Products, www.gembio.com, cat. no. 100-800).

### Cell transformations

Primary porcine pancreatic epithelial cells were grown to 80% confluency in 10 cm dishes under standard conditions, which generally required 14 days. The medium then was exchanged with supernatant from Lenti-X™ 293T cells containing the LV-GK lentiviral particles, along with 1μg/mL polybrene (cat. no. BM-862M, Boston Bioproducts). After 24 h at 37°C, fresh HEK293T supernatant was added to the epithelial cells for a second incubation; after 48 h, the epithelial medium was changed to complete medium and incubated for an additional 24 h. The cells then were split into 6-well plates and cultured under standard conditions 80% confluency. The presence of transduced cells was determined with inverted fluorescent microscopy of living cells.

### Immunoblotting

Immunoblotting was performed to confirm expression of SV40T, decrease or loss of p53, and expression of KRAS^G12D^ (see **Table S3** for a list of antibodies used), as previously described^32^. Immunoblot signal was detected using the Bio-Rad ChemiDoc™ MP Imaging System (bio-rad.com).

### Soft agar assay

A standard soft agar assay^33^ was used to determine anchorage-independent growth. A base layer of 1% agarose was plated into 6-well plates. A total of 20,000 cells/well were mixed with 0.7% agarose (750μL) and plated on top of the base layer. The plates were incubated under standard conditions for 12 days. The cells were then incubated with 200 μL of nitro blue tetrazolium chloride for 24 h to stain the colonies, and then counted using an inverted microscope. Mean count data were obtained from triplicate wells.

### Migration and Invasion assay

Cells were incubated in serum-free medium for 24 h prior to plating the experiment. Basement membrane extract (BME, 1X; 100 μL; cat. no. 3455-096-02, R&D systems, rndsystems.com) was mixed with coating solution (1X; 100μL; cat. no. 3455-096-03, R&D systems) was placed into Falcon™ Cell Culture Inserts (0.4 μm pore size; Falcon cat. no. 353095) and allowed to solidify overnight at 37°C. The next day cells (50,000) were plated into the upper chamber (in triplicate) for both invasion (with BME) or migration analysis (without BME), and incubated under standard conditions for 48 h. The medium from the upper chamber then was aspirated away, and any cells remaining in the upper chamber were removed using a cotton swab. Cells that had migrated to the bottom of the membrane were stained with crystal violet and imaged.

### Population doubling assay

Cells were plated in 6-well plates (20,000 cell/well in 2 mL of complete medium; in triplicate) and cultured under standard conditions. Plates were trypsinized on days 3, 6, 9 and 12, and cells were counted with a hemocytometer. Cell number *vs*. day was plotted to determine the day range of linear growth. These data were used to determine population doubling time (DT) using the formula: DT = (Δt) × ln(2) ÷ ln(N_*f*_/N_*i*_) where Δt = time interval between initial and final cell count, N_*f*_ = cell count at final time, and N_*i*_ = cell count at initial time^34^.

### Proliferation assay

Relative cell proliferation rates were determined using an MTT (3-(4,5-dimethylthiazol-2-yl)-2,5-diphenyltetrazolium bromide) assay kit (Vybrant™ MTT Cell Proliferation Assay Kit, Invitrogen™, Thermo Fisher Scientific, cat. no. V13154). Cells were plated in triplicate in a 96-well plate (3,000 cell/well in 200 μL of complete medium and cultured under standard conditions for 48 h. MTT reagent then was added to the cells per the manufacturer’s instructions, followed by addition of the solvent solution 3.5 h later. Absorbance was measured with a plate reader 3.5 h after solvent addition. Mean absorbance was normalized to absorbance from wild-type PDECs to calculate fold-difference in proliferation.

### Immunofluorescence and immunohistochemistry

Antibodies used in immunofluorescent and immunohistochemical experiments are listed in **Table S3**. Vector Laboratories (vectorlabs.com) ImmPress® goat (MP-7452) or rabbit (MP-7451) anti-mouse IgG polymer kits were used for IHC analyses per the manufacturer’s instructions. Detection of the HRP/ peroxidase enzyme was performed with SignalStain® DAB Substrate Kit from Cell Signaling (cat. no. 8059; www.cellsignal.com), per the manufacturer’s instructions.

### Subcutaneous tumorigenic cell injection

Subcutaneous implantation of tumorigenic cells was performed as previously described^35^, with some modifications. Transformed PDECs were trypsinized, counted, and resuspended in DMEM at a concentration of 5 × 10^6^ cells/mL. Nude mice (N = 10 per treatment group; 100% female; maintained in microisolator cages with soft bedding and fed regular chow ad lib) were allocated into treatment groups using an online randomization tool (www.randomizer.org). Using a 1 mL syringe with a 27 gauge needle, each subject was injected with a single dose of 500,000 cells in 100 μL of DMEM, into the subcutaneous space of the right hind flank without anesthetic. Tumors were allowed to grow for 4 weeks or until they reached 2 cm in diameter, as measured externally with a caliper, and then subjects were euthanized using an AVMA-approved^29^ method of CO_2_ asphyxiation. At necropsy all gross tumor was measured and collected, with samples undergoing formalin fixation, paraffin embedding, and H&E or immunohistochemical staining as described above. Sections were analyzed by an independent, unbiased (blinded to section identify) pathologist to determine tissue of origin and malignant features.

### Statistics and power analysis

Data are reported as mean ± standard deviation. Groups of continuous data were compared with ANOVA and the unpaired t-test. Categorical data were compared with the Fisher or Chi square test. For the power analysis of the murine subcutaneous tumor implant assay, tumor diameter was selected as the endpoint. Setting alpha to 0.05 and power (1 – beta) to 0.8, and with an estimated standard deviation of 20% of the mean, ten mice per group were needed across all treatment groups to detect a 30% difference in mean tumor diameter.

## Results

### Isolation of primary porcine PDECs

Cells cultured from microdissected porcine pancreatic ducts displayed epithelial morphology under phase microscopy (not shown) and stained for CK19 and Pan-Keratin (markers of pancreatic ductal epithelium^36^; **Fig. 1A**). Based on these results, we were confident that we had cultures of primary porcine pancreatic ductal epithelial cells (PDECs) that we could transform into tumorigenic epithelial cell lines.

**Fig. 1.**
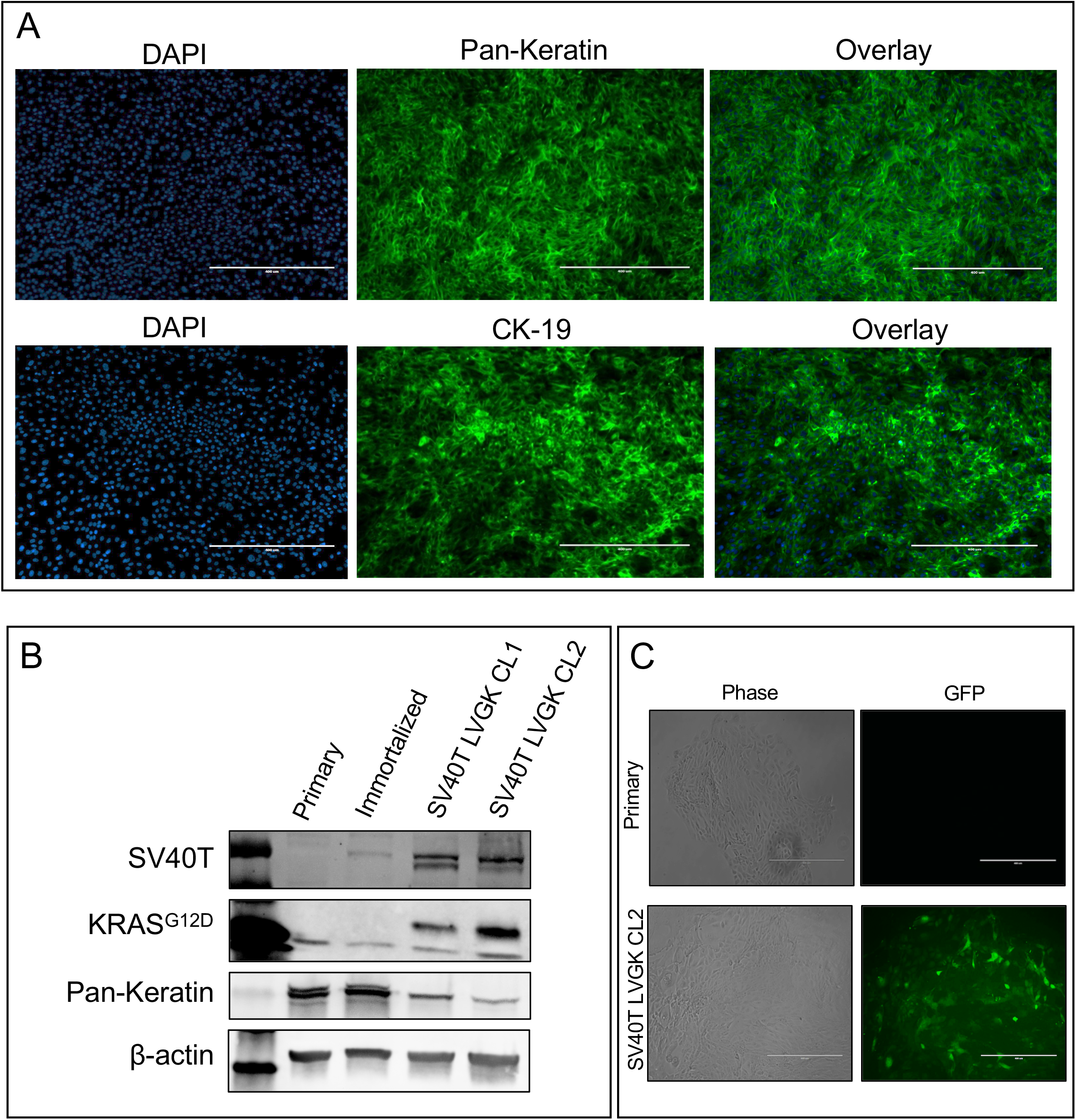
Isolation and transduction of primary porcine pancreatic ductal epithelial cells. (A) Immunofluorescent staining for Pan-Keratin and cytokeratin 19 (CK-19) in cultured primary epithelial cells. (B) Immunoblot for SV40T, mutant KRAS^G12D^, and Pan-Keratin in primary epithelial cells, SV40T immortalized cells, and SV40T immortalized cells transformed with KRAS^G12D^ (SV40T LV-GK; two lines, CL1 and CL2). (C) Expression of ZsGreen1 (a green fluorescent protein variant) in primary epithelial cells and SV40T immortalized cells transduced with the LV-GK; phase and inverted fluorescent microscopy of living cells. Measure bars = 400 μm.

### Generation of transformed PDEC lines

In order to transform our primary PDECs, we first immortalized the cells with SV40 large T antigen (SV40T) to inhibit endogenous p53. Inactivating mutations of the tumor suppressor p53 protein occur in approximately 50-75% of PDAC cases^37–39^. SV40T binds to the same domain of p53 where missense mutations occur, and mimics the effect seen with these mutations^40,41^. Expression of SV40T after lentiviral insertion was confirmed with immunoblotting (**Fig. 1B**). We then generated a lentiviral vector containing the porcine *KRAS*^G12D^ sequence previously identified^30^ as the porcine equivalent to the mutant *KRAS* which is present in multiple human cancers^38,42–45^; expression of the murine version of this mutant KRAS was the basis for the KRAS/p53 (KPC) mouse, a genetically engineered murine model of pancreatic cancer^46^. We utilized a vector (**Fig. S3**) that contained the human cytomegalovirus immediate early promoter that is upstream from a multiple cloning site, which allows for both the gene of interest and a fluorescent protein marker (ZsGreen1^47^) to be translated in a single bicistronic mRNA. Expression of KRAS^G12D^ after lentiviral insertion was confirmed with immunoblotting and fluorescent microscopy of living cells (**Fig. 1B,C**).

### *In vitro* behavior of transformed PDEC lines

Two lines of transformed PDECs (SV40T LV-GK CL1 and SV40T LV-GK CL2, abbreviated as CL1 and CL2) were tested in a series of *in vitro* transformation assays. Both CL1 and CL2 demonstrated increased (≥10-fold) soft agar colony formation over wild type and SV40T immortalized cells (**Fig. 2A)**. Population doubling time was shorter and cellular proliferation (metabolic dye conversion) was greater for both CL1 and CL2 compared to wild type and SV40T cells (**Fig. 2B,C**). Calculation from the data in **Fig. 2B** demonstrated that the doubling time for the transformed cell lines was approximately the same at ~15 h, compared to the ~3 d doubling time of wild type and SV40T cells (a ~5-fold difference). Both CL1 and CL2 had increased migration and invasion capability compared to wild type and SV40T cells (**Fig. 2D,E**). Based on these *in vitro* assays of transformation, we suspected that both our transformed cell lines had the potential to form tumors *in vivo*. With the exception of higher soft agar colony formation in CL2 (**Fig. 2A**), we were unable demonstrate an obvious difference in transformed behavior between CL1 and CL2. Based on the difference in the soft agar colony assay, we elected to utilize CL2 in the below *in vivo* tumorigenicity assay.

**Fig. 2.**
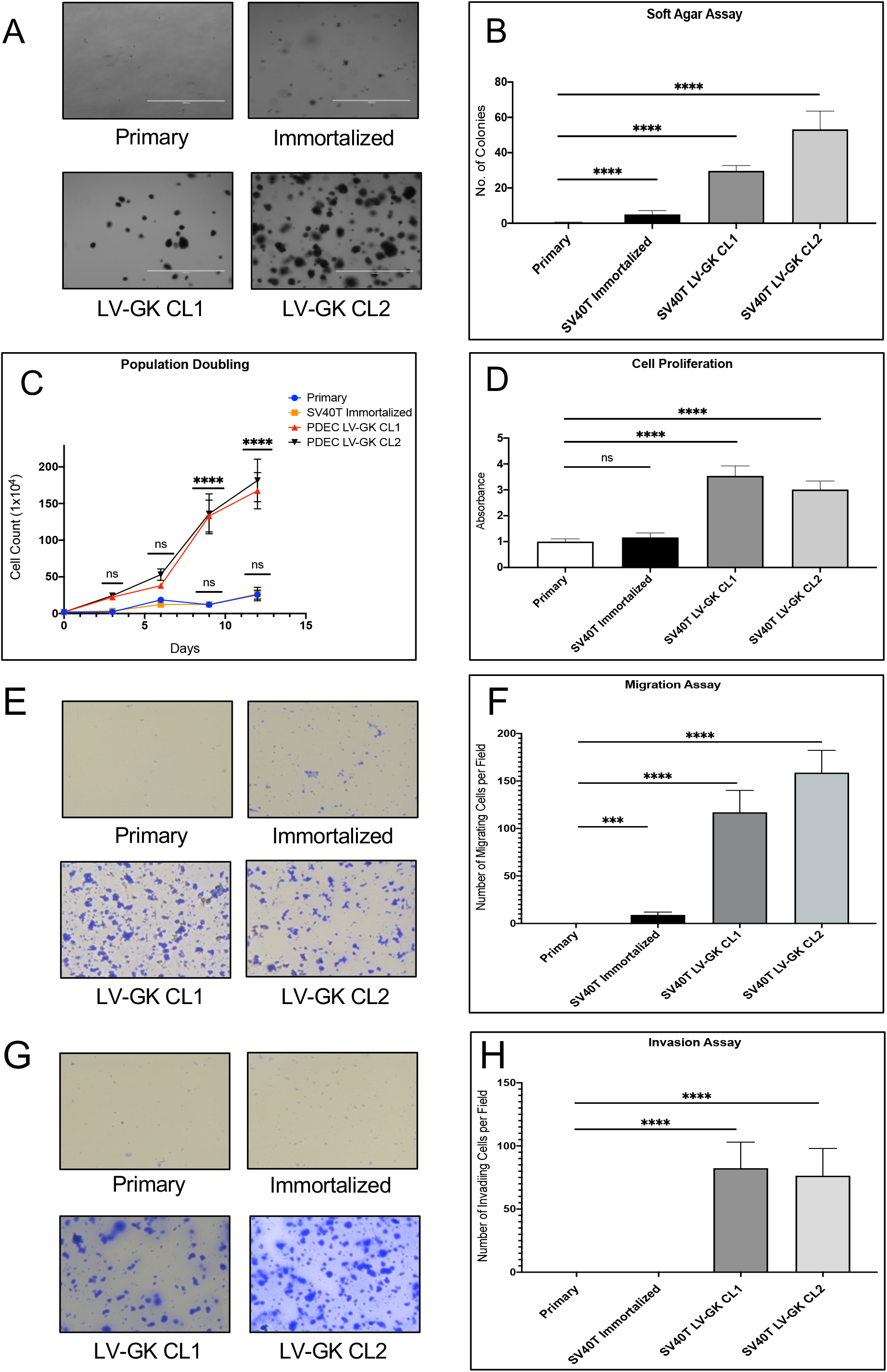
*In vitro* cell transformation assays. Cultured primary epithelial cells from porcine pancreatic duct were compared with SV40T immortalized and transformed (SV40T + KRAS^G12D^) cell lines. (A) Phase contrast images of soft agar assay (bar = 1,000 μm). (B) Plot of soft agar assay. (C) Cell culture population doubling time (count-based analysis). (D) Proliferation rate using metabolic dye assay, represented as fold change with respect to primary cells (defined as 1). (E-F) Transwell migration assay (absence of BME), phase images and plot. (G-H) Transwell invasion assay (presence of BME), phase images and plot. Each bar or data point in this Figure represents the mean ± SD of triplicate wells; each experiment performed three times on separate days (one representative experiment shown in each panel); ****p < 0.0001; ns = not significant (p ≥ 0.05).

### Transformed PDECs generated subcutaneous tumors with PDAC features when injected into immunodeficient mice

In order to determine if transformed PDECs were tumorigenic *in vivo*, we utilized homozygous athymic model of subcutaneous cell implantation (xenografting into nude mice)^16^. Wild type, immortalized SV40T and transformed SV40T LV-GK CL2 cells were injected into the right flank of nude mice (500K cells/site; 1 site/mouse). Neither the wild type nor the immortalized SV40T cells grew tumors. The transformed cell line grew sizeable tumors (~150 mm^3^) within 4 weeks (**Fig. 3A,B**). The subcutaneous tumors were vascularized and mucinous in gross appearance (**Fig. 3A**). Pathological review of H&E staining from these tumor xenografts demonstrated formation of numerous, variably-sized ductal structures surrounded by enlarged atypical nuclei with prominent nucleoli, a few pyknotic nuclei, and some tumor-associated stroma (**Fig. 3C-E**), characteristic of a ductular adenocarcinoma^48^.

**Fig. 3.**
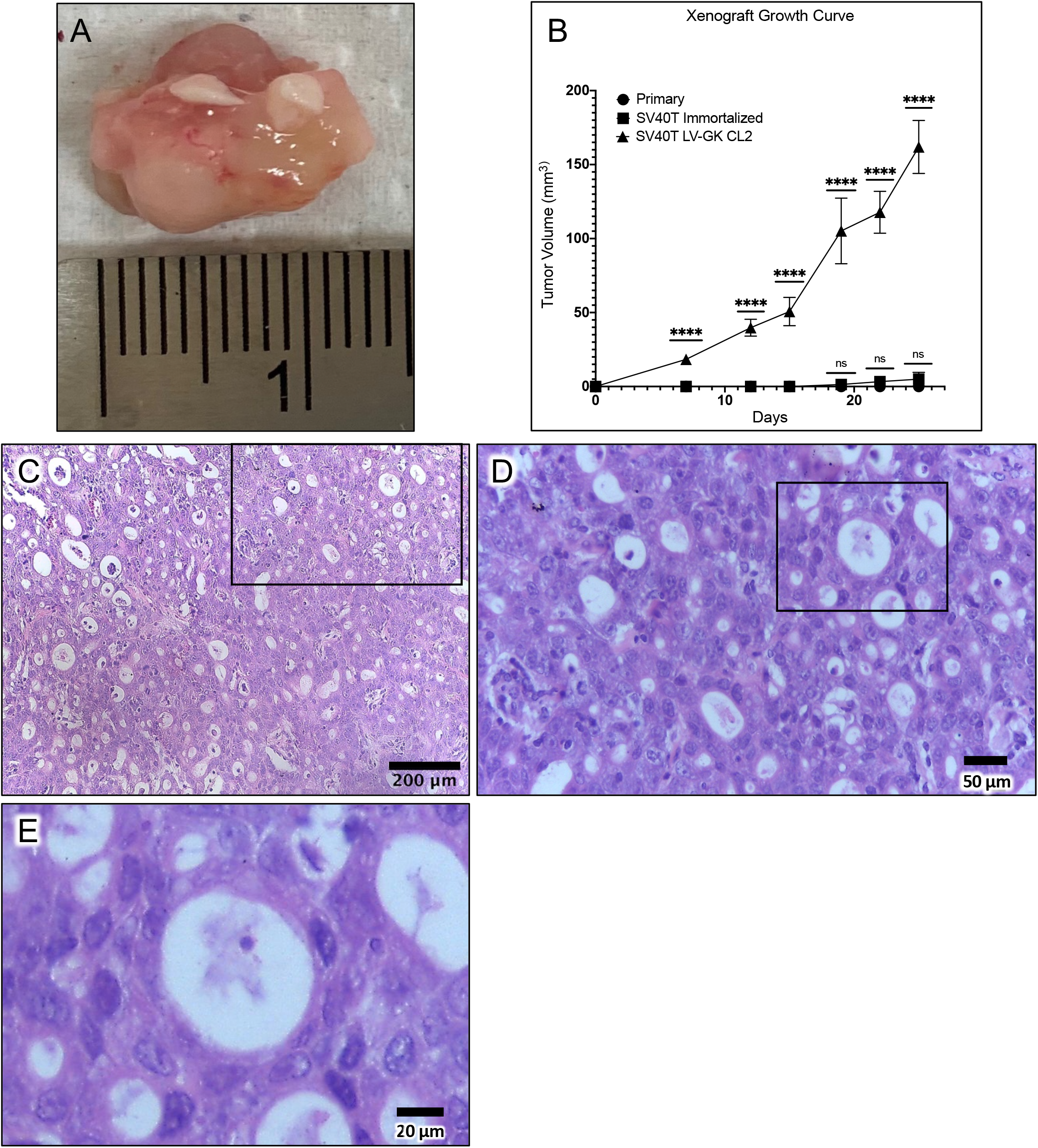
*In vivo* tumorigenesis assay. (A) Representative xenograft tumor explanted from a nude mouse 42 days after subcutaneous injection of transformed epithelial cells (SV40T LV-GK CL2). (B) Comparison of xenograft tumor volume between primary epithelial cells versus the SV40T immortalized and SV40T LV-GK CL2 cell lines; ****p < 0.0001; ns = not significant (p ≥ 0.05). (C) H&E microscopic image of xenograft tumor. (D) Higher power H&E image; region of interest box shown in panel C. (E) Higher power H&E image; region of interest box shown in panel D.

### Immunohistochemical features of tumor xenografts

Tumors from the subcutaneous implant model underwent immunohistochemical staining with Pan-Keratin, CK19, SV40T, KRAS^G12D^, p21, and PCNA (**Fig. 4**). Tumors stained positive for both Pan-Keratin and CK19 (latter not shown). All tumors formed ductal and acinar-like structures (similar to what is seen in human PDAC; **Fig. 4A-C**), consistent with a tumor of ductal and/or alveolar origin. Confirmation of lentiviral transduction was demonstrated with KRAS^G12D^ and SV40T staining (**Fig. 4D-E**). These tumors also stained for PCNA (**Fig. 4F**), a proliferative marker that is expressed in human PDAC and correlates with a poor prognosis^49^. We also stained for p21 to confirm loss of wild type p53, since p21 is activated downstream of p53^50^. Tumor cells did not stain for p21 expression, with the exception of some infiltrating lymphocytes, consistent with loss of p53 function with SV40T expression in the tumor cells (**Fig. 4G**). Control staining for p21 in the skin of these immunodeficient mice is shown in **Fig. S4**.

**Fig. 4.**
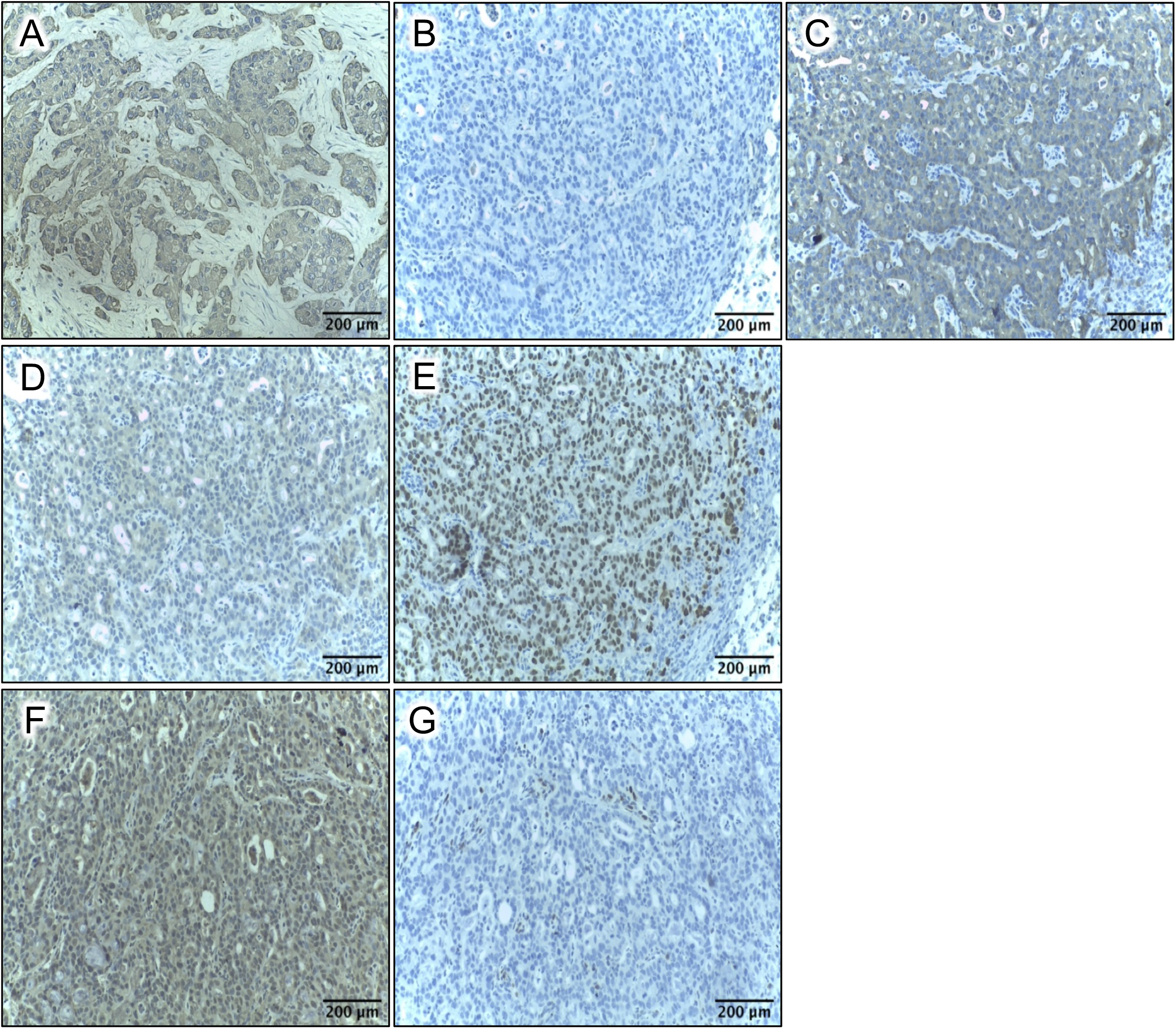
Immunohistochemistry of tumor xenografts from subcutaneous nude mouse assay. (A) Pan-Keratin; section from human PDAC (original tumor, not a xenograft); utilized as a positive control (nuclear counterstain = hematoxylin). Panels B-G are all from xenograft tumor, obtained with implantation of the SV40T LV-GK CL2 cell line. (B) Negative control (all reagents except no primary antibody). (C) Pan-Keratin. (D) KRAS^G12D^ (antibody specific to mutant KRAS). (E) SV40T. (F) PCNA. (G) p21.

## Discussion

Porcine biomedical models have been used for decades in the fields of trauma and hemostasis^51^, xenotransplantation^52,53^, dermal healing^54^, toxicology^55^, atherosclerosis^56^, and cardiac regeneration^57^; the utility of these models is growing. A porcine genome map was generated in 2012^58^, and further coverage, annotation, and confirmation is ongoing^59,60^. Porcine-centered online tools and databases are now available^61^. Genetic manipulation of pigs (including knockouts, tissue-specific transgenics, inducible expression^30,62–69^) with similar tools as used in the mouse is becoming more routine, with new gene-edited porcine models emerging for diseases such as atherosclerosis, cystic fibrosis, Duchenne muscular dystrophy, and ataxia telangiectasia^70,71^.

The rationale to build a porcine model of pancreatic cancer is (1) to have a platform for diagnostic/therapeutic device development otherwise not achievable in murine models; and (2) to have a highly predictive preclinical model in which anti-cancer therapies (including immunotherapies) could be vetted/optimized prior to a clinical trial^72^. The rationale to use the pig in this modeling effort is that this species mimics human genomics^59,73–76^, epigenetics^77^, physiology^55,73,78,79^, metabolism^73,79,80^, inflammation and immune response^76,81–85^, and size^79,86^ remarkably well (in particular, better than mice), with reasonable compromises towards cost and husbandry^79^. So, based on the pig’s relatively large size and its proven track record in replicating human biology which, incidentally, is a demonstrably better replication than can be obtained with rodents, we selected swine as the model organism for this pancreatic cancer project.

Research on immunocompetent large animal cancer models^87–89^ includes prostate cancer, for which there is a canine model^90^. In addition, Munich investigators reported the engineering of (i) an *APC* mutant pig that developed rectal polyposis^17,91^ and (ii) a pig with Cre-inducible p53 deficiency^67^. This group subsequently determined that their p53-null subjects (*TP53*^R167H/R167H^) developed osteosarcoma by age 7-8 months^92^. Other p53-deficient pigs have been engineered since this initial report^68,93^; in the report from Iowa, half (5 out of 10) of p53-deficient (*TP53*^R167H/R167H^) pigs developed lymphoma or osteogenic tumor at age 6-18 months^68^. A group in Denmark reported the creation of a *BRCA* mutant pig in 2012.^94^. In 2017, another genetic porcine model of intestinal neoplasia was reported^95^, utilizing inducible expression of KRAS^G12D^, c-Myc, SV40 large T antigen, and retinoblastoma protein (pRb).

A KRAS/p53 “Oncopig” was reported in 2015^31,88,96^. This subject has a somatic LSL-cassette that can express dominant negative p53 (R167H mutation) and activated KRAS (G12D mutation); i.e., the porcine analog of the KRAS/p53 mouse^46,97^. Site-specific expression of Cre recombinase in the Oncopig resulted in localized p53 inhibition and KRAS activation; subcutaneous injection of AdCre produced mesenchymal tumors at the injection sites^31^ in non-immunosuppressed Oncopigs.

In 2018, induction of autochthonous pancreatic tumor in one KRAS/p53 Oncopig was accomplished^98^ by injection of adenovirus-expressing-Cre into the pancreatic duct, in order to minimize transformation of non-epithelial cells. At the 12-month time point in this single subject, pancreatic tumor was not evident radiographically nor grossly, but was visible after organ sectioning. We recently posted our preliminary data on the induction of pancreatic cancer in 14 Oncopig subjects^99^, in which we observed fulminant, grossly evident pancreatic tumor growth in 10 subjects within 2 weeks of AdCre injection into the duct a surgically-isolated pancreatic lobe. In 2017, initial work was published on a Oncopig-based model of hepatocellular carcinoma^100^.

Löhr et al.^101^ demonstrated that bovine pancreatic ductal cells could be transformed with SV40T and mutant KRAS. They orthotopically implanted these transformed cells into nude mice, and observed pancreatic tumor development and metastasis to the liver. Adam et al.^30^ transformed primary porcine dermal fibroblasts with retroviral insertion of hTERT, p53^DD^ (dominant negative), cyclin D1, CDK4^R24C^, c-Myc^T58A^ and H-Ras^G12V^. These transformed cells also grew tumors after subcutaneous injection in both immunodeficient mice and autologous wild type swine. Of note, the latter required immunosuppression with cyclosporine, prednisone, and azathioprine to prevent the host from rejecting the implanted cells. Our work differs from the previous publications in that we created tumorigenic cell lines by transforming ductal epithelial cells obtained from the porcine pancreas which, similar to bovine pancreatic ductal cells, only required KRAS activation and p53 inhibition.

Our overall goal with this project was to generate tumorigenic pancreatic cell lines that could be used in an immunocompetent porcine model of pancreatic cancer. Murine models can be limited in their ability to replicate human biology and size, so a large animal model of pancreatic cancer likely would enhance our ability to develop and test new diagnostic and treatment modalities for this disease. The data presented herein demonstrated that wild type porcine pancreatic ductal epithelium can be transformed with SV40T and mutant KRAS^G12D^; these transformed cells subsequently can grow tumors in immunodeficient mice, displaying histological features similar to human PDAC. These data provide a pathway for the construction of an orthotopic porcine model of pancreatic cancer, namely, implantation of tumorigenic pancreatic epithelial cells into the pancreas. Future work will investigate the graft-host relationship and immune response in both allogeneic and autologous hosts.

## Acknowledgements

This study is the result of work supported in part with resources and the use of facilities at the Omaha VA Medical Center (Nebraska-Western Iowa Health Care System). The work was supported by NIH grant 5R01CA222907 (MAC). Portions of this study were presented at the 108th Annual Meeting of the American Association for Cancer Research (AACR) in Washington, D.C., April 1-5, 2017. The authors would like to acknowledge the technical assistance of Gerri Siford, Chris Hansen, Kelly O’Connell, and Tom Caffrey. The authors also would like to thank Dr. Lawrence B. Schook and Dr. Laurie A. Rund at the University of Illinois for the gift of their plasmid that contained the *KRAS*^G12D^ mutation, along with comments, insights and suggestions for its use in this project.

## Data Availability Statement

Datasets from this manuscript will be freely shared upon request to the senior author (MAC).

## Author Contributions

| Contributor Role | Description | Author |
| --- | --- | --- |
| Conceptualization | Ideas; formulation or evolution of overarching research goals and aims. | MAH, MAC |
| Data Curation | Management activities to annotate (produce metadata), scrub data and maintain research data (including software code, where it is necessary for interpreting the data itself) for initial use and later reuse. | KB, SBC, NSP, NR |
| Formal Analysis | Application of statistical, mathematical, computational, or other formal techniques to analyze or synthesize study data. | KB, SBC, NSP, NR |
| Funding Acquisition | Acquisition of the financial support for the project leading to this publication. | MAC |
| Investigation | Conducting a research and investigation process, specifically performing the experiments, or data/evidence collection. | KB, SBC, NSP, NR |
| Methodology | Development or design of methodology; creation of models. | KB, SBC, NSP, NR, MAC |
| Project Administration | Management and coordination responsibility for the research activity planning and execution. | MAC |
| Resources | Provision of study materials, reagents, materials, patients, laboratory samples, animals, instrumentation, computing resources, or other analysis tools. | MAH, MAC |
| Software | Programming, software development; designing computer programs; implementation of the computer code and supporting algorithms; testing of existing code components. | na |
| Supervision | Oversight and leadership responsibility for the research activity planning and execution, including mentorship external to the core team. | MAH, MAC |
| Validation | Verification, whether as a part of the activity or separate, of the overall replication/reproducibility of results/experiments and other research outputs. | KB |
| Visualization | Preparation, creation and/or presentation of the published work, specifically visualization/data presentation. | KB, NR |
| Draft Preparation | Creation and/or presentation of the published work, specifically writing the initial draft (including substantive translation). | KB, MAC |
| Review & Editing | Preparation, creation and/or presentation of the published work by those from the original research group, specifically critical review, commentary or revision – including pre- or post-publication stages. | All |

na = not applicable; All = all authors.

## Supporting Information

**Fig. S1**. Responses to ARRIVE Guidelines.

**Fig. S2**. Isolation of Epithelial Cells from Porcine Pancreas.

**Fig. S3**. Vector map.

**Fig. S4**. p21 immunohistochemistry in murine skin.

**Table S1**. Responses to the NIH Preclinical Research Guidelines

**Table S2**. Primers and other short sequences

**Table S3**. Antibody information

## References

1. Saad AM, Turk T, Al-Husseini MJ, Abdel-Rahman O. Trends in pancreatic adenocarcinoma incidence and mortality in the United States in the last four decades; a SEER-based study. BMC Cancer. 2018;18(1):688. PMID: 29940910. DOI: 10.1186/s12885-018-4610-4

2. American Cancer Society. Cancer Facts & Figures 2016. Atlanta: American Cancer Society; 2016. Available from: http://www.cancer.org/research/cancerfactsstatistics/cancerfactsfigures2016/index.

3. SEER. Surveillance, Epidemiology, and End Results Program Stat Fact Sheets: Pancreas Cancer. Accessed December 12, 2016. Available from: URL: http://seer.cancer.gov/statfacts/html/pancreas.html.

4. National Comprehensive Cancer Network. Pancreatic Adenocarcinoma, Version 1.2021. 23 October 2020. In: NCCN Clinical Practice Guidelines In Oncology (NCCN Guidelines®) [Internet]. Available from: https://www.nccn.org.

5. Seok J, Warren HS, Cuenca AG, Mindrinos MN, Baker HV, Xu W, Richards DR, McDonald-Smith GP, Gao H, Hennessy L, Finnerty CC, Lopez CM, Honari S, Moore EE, Minei JP, Cuschieri J, Bankey PE, Johnson JL, Sperry J, Nathens AB, Billiar TR, West MA, Jeschke MG, Klein MB, Gamelli RL, Gibran NS, Brownstein BH, Miller-Graziano C, Calvano SE, Mason PH, Cobb JP, Rahme LG, Lowry SF, Maier RV, Moldawer LL, Herndon DN, Davis RW, Xiao W, Tompkins RG, Inflammation, Host Response to Injury LSCRP. Genomic responses in mouse models poorly mimic human inflammatory diseases. Proc Natl Acad Sci U S A. 2013;110(9):3507–3512. PMID: 23401516. DOI: 10.1073/pnas.1222878110

6. Begley CG, Ellis LM. Drug development: Raise standards for preclinical cancer research. Nature. 2012;483(7391):531–533. PMID: 22460880. DOI: 10.1038/483531a

7. Cook N, Jodrell DI, Tuveson DA. Predictive in vivo animal models and translation to clinical trials. Drug Discov Today. 2012;17(5-6):253–260. PMID: 22493784. DOI: 10.1016/j.drudis.2012.02.003

8. Le Magnen C, Dutta A, Abate-Shen C. Optimizing mouse models for precision cancer prevention. Nat Rev Cancer. 2016;16(3):187–196. PMID: 26893066. DOI: 10.1038/nrc.2016.1

9. Bailey KL, Carlson MA. Porcine Models of Pancreatic Cancer. Front Oncol. 2019;9:144. PMID: 30915276. DOI: 10.3389/fonc.2019.00144

10. Reichert JM, Wenger JB. Development trends for new cancer therapeutics and vaccines. Drug Discov Today. 2008;13(1-2):30–37. PMID: 18190861. DOI: 10.1016/j.drudis.2007.09.003

11. Sharpless NE, Depinho RA. The mighty mouse: genetically engineered mouse models in cancer drug development. Nat Rev Drug Discov. 2006;5(9):741–754. PMID: 16915232. DOI: 10.1038/nrd2110

12. Ebos JM, Kerbel RS. Antiangiogenic therapy: impact on invasion, disease progression, and metastasis. Nat Rev Clin Oncol. 2011;8(4):210–221. PMID: 21364524. DOI: 10.1038/nrclinonc.2011.21

13. Francia G, Cruz-Munoz W, Man S, Xu P, Kerbel RS. Mouse models of advanced spontaneous metastasis for experimental therapeutics. Nat Rev Canc. 2011;11(2):135–141. PMID: 21258397. DOI: doi:10.1038/nrc3001

14. O’Collins VE, Macleod MR, Donnan GA, Horky LL, van der Worp BH, Howells DW. 1,026 experimental treatments in acute stroke. Ann Neurol. 2006;59(3):467–477. PMID: 16453316. DOI: 10.1002/ana.20741

15. Scott S, Kranz JE, Cole J, Lincecum JM, Thompson K, Kelly N, Bostrom A, Theodoss J, Al-Nakhala BM, Vieira FG. Design, power, and interpretation of studies in the standard murine model of ALS. Amyotroph Lat Scler. 2008;9(1):4–15. PMID: 18273714. DOI: 10.1080/17482960701856300

16. Talmadge JE, Singh RK, Fidler IJ, Raz A. Murine models to evaluate novel and conventional therapeutic strategies for cancer. Am J Pathol. 2007;170(3):793–804. PMID: 17322365. DOI: 10.2353/ajpath.2007.060929

17. Flisikowska T, Merkl C, Landmann M, Eser S, Rezaei N, Cui X, Kurome M, Zakhartchenko V, Kessler B, Wieland H, Rottmann O, Schmid RM, Schneider G, Kind A, Wolf E, Saur D, Schnieke A. A porcine model of familial adenomatous polyposis. Gastroenterology. 2012;143(5):1173–1175 e1177. PMID: 22864254. DOI: 10.1053/j.gastro.2012.07.110

18. Rogers CS, Stoltz DA, Meyerholz DK, Ostedgaard LS, Rokhlina T, Taft PJ, Rogan MP, Pezzulo AA, Karp PH, Itani OA, Kabel AC, Wohlford-Lenane CL, Davis GJ, Hanfland RA, Smith TL, Samuel M, Wax D, Murphy CN, Rieke A, Whitworth K, Uc A, Starner TD, Brogden KA, Shilyansky J, McCray PB, Jr., Zabner J, Prather RS, Welsh MJ. Disruption of the CFTR gene produces a model of cystic fibrosis in newborn pigs. Science. 2008;321(5897):1837–1841. PMID: 18818360. DOI: 10.1126/science.1163600

19. Pezzulo AA, Tang XX, Hoegger MJ, Abou Alaiwa MH, Ramachandran S, Moninger TO, Karp PH, Wohlford-Lenane CL, Haagsman HP, van Eijk M, Banfi B, Horswill AR, Stoltz DA, McCray PB, Jr., Welsh MJ, Zabner J. Reduced airway surface pH impairs bacterial killing in the porcine cystic fibrosis lung. Nature. 2012;487(7405):109–113. PMID: 22763554. DOI: 10.1038/nature11130

20. Maddalo D, Manchado E, Concepcion CP, Bonetti C, Vidigal JA, Han YC, Ogrodowski P, Crippa A, Rekhtman N, de Stanchina E, Lowe SW, Ventura A. In vivo engineering of oncogenic chromosomal rearrangements with the CRISPR/Cas9 system. Nature. 2014;516(7531):423–427. PMID: 25337876. DOI: 10.1038/nature13902

21. Sanchez-Rivera FJ, Papagiannakopoulos T, Romero R, Tammela T, Bauer MR, Bhutkar A, Joshi NS, Subbaraj L, Bronson RT, Xue W, Jacks T. Rapid modelling of cooperating genetic events in cancer through somatic genome editing. Nature. 2014;516(7531):428–431. PMID: 25337879. DOI: 10.1038/nature13906

22. Day CP, Merlino G, Van Dyke T. Preclinical mouse cancer models: a maze of opportunities and challenges. Cell. 2015;163(1):39–53. PMID: 26406370. DOI: 10.1016/j.cell.2015.08.068

23. Singh M, Murriel CL, Johnson L. Genetically engineered mouse models: closing the gap between preclinical data and trial outcomes. Cancer Res. 2012;72(11):2695–2700. PMID: 22593194. DOI: 10.1158/0008-5472.CAN-11-2786

24. Zitvogel L, Pitt JM, Daillere R, Smyth MJ, Kroemer G. Mouse models in oncoimmunology. Nat Rev Cancer. 2016;16(12):759–773. PMID: 27687979. DOI: 10.1038/nrc.2016.91

25. Kilkenny C, Browne WJ, Cuthill IC, Emerson M, Altman DG. Improving bioscience research reporting: the ARRIVE guidelines for reporting animal research. PLoS Biol. 2010;8(6):e1000412. PMID: 20613859. DOI: 10.1371/journal.pbio.1000412

26. National Institutes of Health. Principles and Guidelines for Reporting Preclinical Research. 17 December 2017. Available from: https://www.nih.gov/research-training/rigor-reproducibility/principles-guidelines-reporting-preclinical-research.

27. National Institutes of Health. Enhancing Reproducibility through Rigor and Transparency. June 9, 2015. http://grants.nih.gov/grants/guide/notice-files/NOT-OD-15-103.html

28. Committee for the Update of the Guide for the Care and Use of Laboratory Animals. Guide for the Care and Use of Laboratory AnimalsWashington, DC: The National Academies Press; 2011.

29. American Veterinary Medical Association Panel on Euthanasia. AVMA Guidelines for the Euthanasia of Animals: 2013 EditionSchaumberg, IL: American Veterinary Medical Association; 2013.

30. Adam SJ, Rund LA, Kuzmuk KN, Zachary JF, Schook LB, Counter CM. Genetic induction of tumorigenesis in swine. Oncogene. 2007;26(7):1038–1045. PMID: 16964292. DOI: 10.1038/sj.onc.1209892

31. Schook LB, Collares TV, Hu W, Liang Y, Rodrigues FM, Rund LA, Schachtschneider KM, Seixas FK, Singh K, Wells KD, Walters EM, Prather RS, Counter CM. A Genetic Porcine Model of Cancer. PLOS ONE. 2015;10(7):e0128864. PMID: 26132737. DOI: 10.1371/journal.pone.0128864

32. Grinnell F, Zhu M, Carlson MA, Abrams JM. Release of mechanical tension triggers apoptosis of human fibroblasts in a model of regressing granulation tissue. Exp Cell Res. 1999;248(2):608–619. PMID: 10222153. DOI: 10.1006/excr.1999.4440

33. Macpherson I, Montagnier L. Agar Suspension Culture for the Selective Assay of Cells Transformed by Polyoma Virus. Virology. 1964;23:291–294. PMID: 14187925. DOI:

34. Mehrara E, Forssell-Aronsson E, Ahlman H, Bernhardt P. Specific growth rate versus doubling time for quantitative characterization of tumor growth rate. Cancer Res. 2007;67(8):3970–3975. PMID: 17440113. DOI: 10.1158/0008-5472.CAN-06-3822

35. Tsutsumida H, Swanson BJ, Singh PK, Caffrey TC, Kitajima S, Goto M, Yonezawa S, Hollingsworth MA. RNA interference suppression of MUC1 reduces the growth rate and metastatic phenotype of human pancreatic cancer cells. Clin Cancer Res. 2006;12(10):2976–2987. PMID: 16707592. DOI: 10.1158/1078-0432.CCR-05-1197

36. Gioviale MC, Damiano G, Montalto G, Buscemi G, Romano M, Lo Monte AI. Isolation and culture of beta-like cells from porcine Wirsung duct. Transplant Proc. 2009;41(4):1363–1366. PMID: 19460560. DOI: 10.1016/j.transproceed.2009.02.062

37. Corbo V, Mafficini A, Amato E, Scarpa A. Pancreatic Cancer Genomics. Cancer Genomics: Springer, 2013, pp. 219–253. ISBN: 9400758413.

38. Waddell N, Pajic M, Patch AM, Chang DK, Kassahn KS, Bailey P, Johns AL, Miller D, al. e. Whole genomes redefine the mutational landscape of pancreatic cancer. Nature. 2015;518:495–501. PMID: PMC4523082. DOI: 10.1038/nature14169

39. Bailey P, Chang DK, Nones K, Johns AL, Patch AM, Gingras MC, Miller DK, Christ AN, Bruxner TJ, Quinn MC, Nourse C, Murtaugh LC, Harliwong I, Idrisoglu S, Manning S, Nourbakhsh E, Wani S, Fink L, Holmes O, Chin V, Anderson MJ, Kazakoff S, Leonard C, Newell F, Waddell N, Wood S, Xu Q, Wilson PJ, Cloonan N, Kassahn KS, Taylor D, Quek K, Robertson A, Pantano L, Mincarelli L, Sanchez LN, Evers L, Wu J, Pinese M, Cowley MJ, Jones MD, Colvin EK, Nagrial AM, Humphrey ES, Chantrill LA, Mawson A, Humphris J, Chou A, Pajic M, Scarlett CJ, Pinho AV, Giry-Laterriere M, Rooman I, Samra JS, Kench JG, Lovell JA, Merrett ND, Toon CW, Epari K, Nguyen NQ, Barbour A, Zeps N, Moran-Jones K, Jamieson NB, Graham JS, Duthie F, Oien K, Hair J, Grutzmann R, Maitra A, Iacobuzio-Donahue CA, Wolfgang CL, Morgan RA, Lawlor RT, Corbo V, Bassi C, Rusev B, Capelli P, Salvia R, Tortora G, Mukhopadhyay D, Petersen GM, Australian Pancreatic Cancer Genome I, Munzy DM, Fisher WE, Karim SA, Eshleman JR, Hruban RH, Pilarsky C, Morton JP, Sansom OJ, Scarpa A, Musgrove EA, Bailey UM, Hofmann O, Sutherland RL, Wheeler DA, Gill AJ, Gibbs RA, Pearson JV, Waddell N, Biankin AV, Grimmond SM. Genomic analyses identify molecular subtypes of pancreatic cancer. Nature. 2016;531(7592):47–52. PMID: 26909576. DOI: 10.1038/nature16965

40. Bargonetti J, Reynisdottir I, Friedman PN, Prives C. Site-specific binding of wild-type p53 to cellular DNA is inhibited by SV40 T antigen and mutant p53. Genes Dev. 1992;6(10):1886–1898. PMID: 1398068. DOI: 10.1101/gad.6.10.1886

41. Dunne RF, Hezel AF. Genetics and Biology of Pancreatic Ductal Adenocarcinoma. Hematol Oncol Clin North Am. 2015;29(4):595–608. PMID: 26226899. DOI: 10.1016/j.hoc.2015.04.003

42. Govindan R, Ding L, Griffith M, Subramanian J, Dees ND, Kanchi KL, Maher CA, Fulton R, Fulton L, Wallis J, Chen K, Walker J, McDonald S, Bose R, Ornitz D, Xiong D, You M, Dooling DJ, Watson M, Mardis ER, Wilson RK. Genomic landscape of non-small cell lung cancer in smokers and never-smokers. Cell. 2012;150(6):1121–1134. PMID: 22980976. DOI: 10.1016/j.cell.2012.08.024

43. Kandoth C, McLellan MD, Vandin F, Ye K, Niu B, Lu C, Xie M, Zhang Q, McMichael JF, Wyczalkowski MA, Leiserson MD, Miller CA, Welch JS, Walter MJ, Wendl MC, Ley TJ, Wilson RK, Raphael BJ, Ding L. Mutational landscape and significance across 12 major cancer types. Nature. 2013;502(7471):333–339. PMID: 24132290. DOI: 10.1038/nature12634

44. Stephens PJ, Tarpey PS, Davies H, Van Loo P, Greenman C, Wedge DC, Nik-Zainal S, Martin S, Varela I, Bignell GR. The landscape of cancer genes and mutational processes in breast cancer. Nature. 2012;486(7403):400–404. PMID: 22722201. DOI:

45. Vogelstein B, Papadopoulos N, Velculescu VE, Zhou S, Diaz LA, Jr., Kinzler KW. Cancer genome landscapes. Science. 2013;339(6127):1546–1558. PMID: 23539594. DOI: 10.1126/science.1235122

46. Hingorani SR, Wang L, Multani AS, Combs C, Deramaudt TB, Hruban RH, Rustgi AK, Chang S, Tuveson DA. Trp53R172H and KrasG12D cooperate to promote chromosomal instability and widely metastatic pancreatic ductal adenocarcinoma in mice. Canc Cell. 2005;7(5):469–483. PMID: 15894267. DOI: 10.1016/j.ccr.2005.04.023

47. Matz MV, Fradkov AF, Labas YA, Savitsky AP, Zaraisky AG, Markelov ML, Lukyanov SA. Fluorescent proteins from nonbioluminescent Anthozoa species. Nat Biotechnol. 1999;17(10):969–973. PMID: 10504696. DOI: 10.1038/13657

48. Hackeng WM, Hruban RH, Offerhaus GJ, Brosens LA. Surgical and molecular pathology of pancreatic neoplasms. Diagn Pathol. 2016;11(1):47. PMID: 27267993. DOI: 10.1186/s13000-016-0497-z

49. Smith SJ, Li CM, Lingeman RG, Hickey RJ, Liu Y, Malkas LH, Raoof M. Molecular Targeting of Cancer-Associated PCNA Interactions in Pancreatic Ductal Adenocarcinoma Using a Cell-Penetrating Peptide. Mol Ther Oncolytics. 2020;17:250–256. PMID: 32368614. DOI: 10.1016/j.omto.2020.03.025

50. Benson EK, Mungamuri SK, Attie O, Kracikova M, Sachidanandam R, Manfredi JJ, Aaronson SA. p53-dependent gene repression through p21 is mediated by recruitment of E2F4 repression complexes. Oncogene. 2014;33(30):3959–3969. PMID: 24096481. DOI: 10.1038/onc.2013.378

51. Kheirabadi BS, Mace JE, Terrazas IB, Fedyk CG, Estep JS, Dubick MA, Blackbourne LH. Safety evaluation of new hemostatic agents, smectite granules, and kaolin-coated gauze in a vascular injury wound model in swine. J Trauma. 2010;68(2):269–278. PMID: 20154537. DOI: 10.1097/TA.0b013e3181c97ef1

52. Cooper DK, Ekser B, Ramsoondar J, Phelps C, Ayares D. The role of genetically engineered pigs in xenotransplantation research. J Pathol. 2016;238(2):288–299. PMID: 26365762. DOI: 10.1002/path.4635

53. Niu D, Wei HJ, Lin L, George H, Wang T, Lee IH, Zhao HY, Wang Y, Kan Y, Shrock E, Lesha E, Wang G, Luo Y, Qing Y, Jiao D, Zhao H, Zhou X, Wang S, Wei H, Guell M, Church GM, Yang L. Inactivation of porcine endogenous retrovirus in pigs using CRISPR-Cas9. Science. 2017;357(6357):1303–1307. PMID: 28798043. DOI: 10.1126/science.aan4187

54. Sullivan TP, Eaglstein WH, Davis SC, Mertz P. The pig as a model for human wound healing. Wound Repair Regen. 2001;9(2):66–76. PMID: 11350644. DOI: 10.1046/j.1524-475x.2001.00066.x

55. Swindle MM, Makin A, Herron AJ, Clubb FJ, Jr., Frazier KS. Swine as models in biomedical research and toxicology testing. Vet Pathol. 2012;49(2):344–356. PMID: 21441112. DOI: 10.1177/0300985811402846

56. Xiangdong L, Yuanwu L, Hua Z, Liming R, Qiuyan L, Ning L. Animal models for the atherosclerosis research: a review. Protein Cell. 2011;2(3):189–201. PMID: 21468891. DOI: 10.1007/s13238-011-1016-3

57. Gouadon E, Moore-Morris T, Smit NW, Chatenoud L, Coronel R, Harding SE, Jourdon P, Lambert V, Rucker-Martin C, Puceat M. Concise Review: Pluripotent Stem Cell-Derived Cardiac Cells, A Promising Cell Source for Therapy of Heart Failure: Where Do We Stand? Stem Cells. 2016;34(1):34–43. PMID: 26352327. DOI: 10.1002/stem.2205

58. Groenen MA, Archibald AL, Uenishi H, Tuggle CK, Takeuchi Y, Rothschild MF, Rogel-Gaillard C, Park C, Milan D, Megens H-J. Analyses of pig genomes provide insight into porcine demography and evolution. Nature. 2012;491(7424):393–398. PMID: 23151582. DOI: 10.1038/nature11622

59. Walters E, Wolf E, Whyte J, Mao J, Renner S, Nagashima H, Kobayashi E, Zhao J, Wells K, Critser J. Completion of the swine genome will simplify the production of swine as a large animal biomedical model. BMC Med Genom. 2012;5(1):55. PMID: 23151353. DOI: 10.1186/1755-8794-5-55

60. Schook LB, Collares TV, Darfour-Oduro KA, De AK, Rund LA, Schachtschneider KM, Seixas FK. Unraveling the swine genome: implications for human health. Annu Rev Anim Biosci. 2015;3:219–244. PMID: 25689318. DOI: 10.1146/annurev-animal-022114-110815

61. Dawson HD, Chen C, Gaynor B, Shao J, Urban JF, Jr. The porcine translational research database: a manually curated, genomics and proteomics-based research resource. BMC Genomics. 2017;18(1):643. PMID: 28830355. DOI: 10.1186/s12864-017-4009-7

62. Gun G, Kues WA. Current progress of genetically engineered pig models for biomedical research. Biores Open Access. 2014;3(6):255–264. PMID: 25469311. DOI: 10.1089/biores.2014.0039

63. Fan N, Lai L. Genetically modified pig models for human diseases. J Genet Genomics. 2013;40(2):67–73. PMID: 23439405. DOI: 10.1016/j.jgg.2012.07.014

64. Prather RS, Lorson M, Ross JW, Whyte JJ, Walters E. Genetically engineered pig models for human diseases. Annu Rev Anim Biosci. 2013;1:203–219. PMID: 25387017. DOI: 10.1146/annurev-animal-031412-103715

65. Luo Y, Li J, Liu Y, Lin L, Du Y, Li S, Yang H, Vajta G, Callesen H, Bolund L, Sorensen CB. High efficiency of BRCA1 knockout using rAAV-mediated gene targeting: developing a pig model for breast cancer. Transgenic Res. 2011;20(5):975–988. PMID: 21181439. DOI: 10.1007/s11248-010-9472-8

66. Luo Y, Bolund L, Sorensen CB. Pig gene knockout by rAAV-mediated homologous recombination: comparison of BRCA1 gene knockout efficiency in Yucatan and Gottingen fibroblasts with slightly different target sequences. Transgenic Res. 2012;21(3):671–676. PMID: 22020980. DOI: 10.1007/s11248-011-9563-1

67. Leuchs S, Saalfrank A, Merkl C, Flisikowska T, Edlinger M, Durkovic M, Rezaei N, Kurome M, Zakhartchenko V, Kessler B. Inactivation and inducible oncogenic mutation of p53 in gene targeted pigs. PLoS ONE. 2012;7(10):e43323. PMID: 23071491. DOI: 10.1371/journal.pone.0043323

68. Sieren JC, Meyerholz DK, Wang XJ, Davis BT, Newell JD, Jr., Hammond E, Rohret JA, Rohret FA, Struzynski JT, Goeken JA, Naumann PW, Leidinger MR, Taghiyev A, Van Rheeden R, Hagen J, Darbro BW, Quelle DE, Rogers CS. Development and translational imaging of a TP53 porcine tumorigenesis model. J Clin Invest. 2014;124(9):4052–4066. PMID: 25105366. DOI: 10.1172/JCI75447

69. Yang L, Guell M, Niu D, George H, Lesha E, Grishin D, Aach J, Shrock E, Xu W, Poci J, Cortazio R, Wilkinson RA, Fishman JA, Church G. Genome-wide inactivation of porcine endogenous retroviruses (PERVs). Science. 2015;350(6264):1101–1104. PMID: 26456528. DOI: 10.1126/science.aad1191

70. Beraldi R, Meyerholz D, Savinov A, Kovács A, Weimer J, Dykstra J, Geraets R, Pearce D. Genetic Ataxia Telangiectasia porcine model phenocopies the multisystemic features of the human disease. Biochim Biophys Acta. 2017:ePub ahead of print, 2017 Jul 2023. PMID: 28746835. DOI: 10.1016/j.bbadis.2017.07.020

71. Perleberg C, Kind A, Schnieke A. Genetically engineered pigs as models for human disease. Dis Model Mech. 2018;11(1):dmm030783. PMID: 29419487. DOI: 10.1242/dmm.030783

72. Segatto NV, Remiao MH, Schachtschneider KM, Seixas FK, Schook LB, Collares T. The Oncopig Cancer Model as a Complementary Tool for Phenotypic Drug Discovery. Front Pharmacol. 2017;8:894. PMID: 29259556. DOI: 10.3389/fphar.2017.00894

73. Kuzmuk KN, Schook LB. Pigs as a Model for Biomedical Sciences. In: Rothschild MF, Ruvinsky A, editors. The Genetics of the Pig. Oxfordshire, UK: CABI, 2011, pp. 426–444. ISBN: 1845937562.

74. Emes RD, Goodstadt L, Winter EE, Ponting CP. Comparison of the genomes of human and mouse lays the foundation of genome zoology. Hum Mol Genet. 2003;12(7):701–709. PMID: 12651866. DOI: 10.1093/hmg/ddg078

75. Dawson HD, McAnulty P, Dayan A, Ganderup N, Hastings K. A comparative assessment of the pig, mouse and human genomes. In: McAnulty P, Dayan A, Ganderup N, Hastings K, editors. The Minipig in Biomedical Research. Boca Raton, FL: CRC Press, 2012, pp. 323–342. ISBN:

76. Dawson HD, Smith AD, Chen C, Urban JF, Jr. An in-depth comparison of the porcine, murine and human inflammasomes; lessons from the porcine genome and transcriptome. Vet Microbiol. 2017;202:2–15. PMID: 27321134. DOI: 10.1016/j.vetmic.2016.05.013

77. Schachtschneider KM, Madsen O, Park C, Rund LA, Groenen MA, Schook LB. Adult porcine genome-wide DNA methylation patterns support pigs as a biomedical model. BMC Genomics. 2015;16:743. PMID: 26438392. DOI: 10.1186/s12864-015-1938-x

78. Vodicka P, Smetana K, Jr., Dvorankova B, Emerick T, Xu YZ, Ourednik J, Ourednik V, Motlik J. The miniature pig as an animal model in biomedical research. Ann N Y Acad Sci. 2005;1049:161–171. PMID: 15965115. DOI: 10.1196/annals.1334.015

79. Swindle MM, Smith AC. Swine in the Laboratory: Surgery, Anesthesia, Imaging, and Experimental Techniques. 3rd. Boca Raton, FL: CRC Press; 2016.

80. Spurlock ME, Gabler NK. The development of porcine models of obesity and the metabolic syndrome. J Nutr. 2008;138(2):397–402. PMID: 18203910. DOI:

81. Röthkotter HJ. Anatomical particularities of the porcine immune system--a physician’s view. Dev Comp Immunol. 2009;33(3):267–272. PMID: 18775744. DOI: 10.1016/j.dci.2008.06.016

82. Bailey M. The mucosal immune system: recent developments and future directions in the pig. Dev Comp Immunol. 2009;33(3):375–383. PMID: 18760299. DOI: 10.1016/j.dci.2008.07.003

83. Fairbairn L, Kapetanovic R, Sester DP, Hume DA. The mononuclear phagocyte system of the pig as a model for understanding human innate immunity and disease. J Leukoc Biol. 2011;89(6):855–871. PMID: 21233410. DOI: 10.1189/jlb.1110607

84. Meurens F, Summerfield A, Nauwynck H, Saif L, Gerdts V. The pig: a model for human infectious diseases. Trends Microbiol. 2012;20(1):50–57. PMID: 22153753. DOI: 10.1016/j.tim.2011.11.002

85. Petersen B, Carnwath JW, Niemann H. The perspectives for porcine-to-human xenografts. Comp Immunol Microbiol Infect Dis. 2009;32(2):91–105. PMID: 18280567. DOI: 10.1016/j.cimid.2007.11.014

86. Ferrer J, Scott WE, 3rd, Weegman BP, Suszynski TM, Sutherland DE, Hering BJ, Papas KK. Pig pancreas anatomy: implications for pancreas procurement, preservation, and islet isolation. Transplantation. 2008;86(11):1503–1510. PMID: 19077881. DOI: 10.1097/TP.0b013e31818bfda1

87. Flisikowska T, Kind A, Schnieke A. Pigs as models of human cancers. Theriogenology. 2016;86(1):433–437. PMID: 27156684. DOI: 10.1016/j.theriogenology.2016.04.058

88. Schachtschneider KM, Schwind RM, Newson J, Kinachtchouk N, Rizko M, Mendoza-Elias N, Grippo P, Principe DR, Park A, Overgaard NH, Jungersen G, Garcia KD, Maker AV, Rund LA, Ozer H, Gaba RC, Schook LB. The Oncopig Cancer Model: An Innovative Large Animal Translational Oncology Platform. Front Oncol. 2017;7:190. PMID: 28879168. DOI: 10.3389/fonc.2017.00190

89. Watson AL, Carlson DF, Largaespada DA, Hackett PB, Fahrenkrug SC. Engineered Swine Models of Cancer. Front Genet. 2016;7:78. PMID: 27242889. DOI: 10.3389/fgene.2016.00078

90. Ittmann M, Huang J, Radaelli E, Martin P, Signoretti S, Sullivan R, Simons BW, Ward JM, Robinson BD, Chu GC. Animal Models of Human Prostate Cancer: The Consensus Report of the New York Meeting of the Mouse Models of Human Cancers Consortium Prostate Pathology Committee. Canc Res. 2013;73(9):2718–2736. PMID: 23610450. DOI: 10.1158/0008-5472.CAN-12-4213

91. Stachowiak M, Flisikowska T, Bauersachs S, Perleberg C, Pausch H, Switonski M, Kind A, Saur D, Schnieke A, Flisikowski K. Altered microRNA profiles during early colon adenoma progression in a porcine model of familial adenomatous polyposis. Oncotarget. 2017;8(56):96154–96160. PMID: 29221194. DOI: 10.18632/oncotarget.21774

92. Saalfrank A, Janssen KP, Ravon M, Flisikowski K, Eser S, Steiger K, Flisikowska T, Muller-Fliedner P, Schulze E, Bronner C, Gnann A, Kappe E, Bohm B, Schade B, Certa U, Saur D, Esposito I, Kind A, Schnieke A. A porcine model of osteosarcoma. Oncogenesis. 2016;5:e210. PMID: 26974205. DOI: 10.1038/oncsis.2016.19

93. Shen Y, Xu K, Yuan Z, Guo J, Zhao H, Zhang X, Zhao L, Qing Y, Li H, Pan W, Jia B, Zhao HY, Wei HJ. Efficient generation of P53 biallelic knockout Diannan miniature pigs via TALENs and somatic cell nuclear transfer. J Transl Med. 2017;15(1):224. PMID: 29100547. DOI: 10.1186/s12967-017-1327-0

94. Luo Y, Kofod-Olsen E, Christensen R, Sorensen CB, Bolund L. Targeted genome editing by recombinant adeno-associated virus (rAAV) vectors for generating genetically modified pigs. J Gen Genom. 2012;39(6):269–274. PMID: 22749014. DOI: 10.1016/j.jgg.2012.05.004

95. Callesen MM, Arnadottir SS, Lyskjaer I, Orntoft MW, Hoyer S, Dagnaes-Hansen F, Liu Y, Li R, Callesen H, Rasmussen MH, Berthelsen MF, Thomsen MK, Schweiger PJ, Jensen KB, Laurberg S, Orntoft TF, Elverlov-Jakobsen JE, Andersen CL. A genetically inducible porcine model of intestinal cancer. Mol Oncol. 2017;11(11):1616–1629. PMID: 28881081. DOI: 10.1002/1878-0261.12136

96. Rodrigues F, Hu W, Rund L, Liang Y, Counter C, Schook L. Development of the “Onco-Pig”: An Inducible Transgenic Porcine Model for Human Cancer [poster]. 2013 Brazilian Genetics Conference; URL: http://www.dbs.illinois.edu/comparativegenomics/userfiles/file/ocnopigPoster_Brazilian%20Genetics%20Conference2013.pdf.

97. DuPage M, Dooley AL, Jacks T. Conditional mouse lung cancer models using adenoviral or lentiviral delivery of Cre recombinase. Nat Protoc. 2009;4(7):1064–1072. PMID: 19561589. DOI: 10.1038/nprot.2009.95

98. Principe DR, Overgaard NH, Park AJ, Diaz AM, Torres C, McKinney R, Dorman MJ, Castellanos K, Schwind R, Dawson DW, Rana A, Maker A, Munshi HG, Rund LA, Grippo PJ, Schook LB. KRAS(G12D) and TP53(R167H) Cooperate to Induce Pancreatic Ductal Adenocarcinoma in Sus scrofa Pigs. Sci Rep. 2018;8(1):12548. PMID: 30135483. DOI: 10.1038/s41598-018-30916-6

99. Patel NS, Bailey K, Lazenby AJ, Carlson MA. Induction of pancreatic neoplasia in the KRAS/TP53 Oncopig: preliminary report. bioRxiv. 2020; published online 2 June 2020. PMID: DOI: 10.1101/2020.05.29.123547

100. Schachtschneider K, Schwind R, Darfour-Oduro K, De A, Rund L, Singh K, Principe D, Guzman G, Ray Jr C, Ozer H. A validated, transitional and translational porcine model of hepatocellular carcinoma. Oncotarget. 2017:Epub ahead of print, 2017 Jun 2029. PMID: 28733541. DOI: 10.18632/oncotarget.18872

101. Lohr M, Muller P, Zauner I, Schmidt C, Trautmann B, Thevenod F, Capella G, Farre A, Liebe S, Jesenofsky R. Immortalized bovine pancreatic duct cells become tumorigenic after transfection with mutant k-ras. Virchows Arch. 2001;438(6):581–590. PMID: 11469690. DOI: 10.1007/s004280100397

