## Supplemental Information for "Porcine pancreatic ductal epithelial cells transformed with KRAS^G12D^ and SV40T are tumorigenic"

SUPPORTING INFORMATION

**Table of Contents**

| | Page |
|---|---|
| **Fig. S1**. Responses to ARRIVE Guidelines. | 2 |
| **Fig. S2**. Isolation of Epithelial Cells from Porcine Pancreas. | 5 |
| **Fig. S3**. Vector maps of pLenti-SV40T and pLVX-IRES-ZsGreen1. | 6 |
| **Fig. S4**. p21 immunohistochemistry. | 7 |
| **Table S1**. Responses to the NIH Preclinical Research Guidelines. | 8 |
| **Table S2**. Primers and other short sequences. | 11 |
| **Table S3**. Antibody information. | 12 |

**Fig. S1**. Responses to the ARRIVE Guidelines (*PLoS Biol* 2010;8(6):e1000412. PMID: 20613859).

| Item | Section | Recommendation | Location of Response |
|---|---|---|---|
| 1 | Title | Provide as accurate and concise a description of the content of the article as possible. | Title page |
| 2 | Abstract | Provide an accurate summary of the background, research objectives, including details of the species or strain of animal used, key methods, principal findings and conclusions of the study. | Abstract page |
| 3 | Introduction: Background | (a) Include sufficient scientific background (including relevant references to previous work) to understand the motivation and context for the study, and explain the experimental approach and rationale. | Introduction |
| | | (b) Explain how and why the animal species and model being used can address the scientific objectives and, where appropriate, the study's relevance to human biology. | Introduction |
| 4 | Introduction: Objectives | Clearly describe the primary and any secondary objectives of the study, or specific hypotheses being tested. | Introduction |
| 5 | Methods: Ethical Statement | Indicate the nature of the ethical review permissions, relevant licences (e.g. Animal [Scientific Procedures] Act 1986), and national or institutional guidelines for the care and use of animals, that cover the research. | Methods |
| 6 | Methods: Study Design | For each experiment, give brief details of the study design including:<br>(a) The number of experimental and control groups. | Methods |
| | | (b) Any steps taken to minimise the effects of subjective bias when allocating animals to treatment (e.g. randomisation procedure) and when assessing results(e.g. if done, describe who was blinded and when). | Methods |
| | | (c) The experimental unit (e.g. a single animal, group or cage of animals). A time-line diagram or flow chart can be useful to illustrate how complex study designs were carried out. | Methods |
| 7 | Methods: Experimental Procedures | For each experiment and each experimental group, including controls, provide precise details of all procedures carried out.<br><br>For example: | Methods |
| | | (a) How (e.g. drug formulation and dose, site and route of administration, anaesthesia and analgesia used [including monitoring], surgical procedure, method of euthanasia). Provide details of any specialist equipment used, including supplier(s).<br>(b) When (e.g. time of day). | Methods |

| | | | |
|---|---|---|---|
| | | (c) Where (e.g. home cage, laboratory, water maze).<br><br>(d) Why (e.g. rationale for choice of specific anaesthetic, route of administration, drug dose used). | |
| 8 | Methods: Experimental Animals | (a) Provide details of the animals used, including species, strain, sex, developmental stage (e.g. mean or median age plus age range) and weight (e.g. mean or median weight plus weight range). | Methods |
| | | (b) Provide further relevant information such as the source of animals, international strain nomenclature, genetic modification status (e.g. knock-out or transgenic), genotype, health/immune status, drug or test naïve, previous procedures, etc. | Methods |
| 9 | Methods: Housing and Husbandry | Provide details of: | |
| | | (a) Housing (type of facility e.g. specific pathogen free [SPF]; type of cage or housing; bedding material; number of cage companions; tank shape and material etc. for fish). | Methods |
| | | (b) Husbandry conditions (e.g. breeding programme, light/dark cycle, temperature, quality of water etc for fish, type of food, access to food and water, environmental enrichment). | Methods |
| | | (c) Welfare-related assessments and interventions that were carried out prior to, during, or after the experiment. | Methods |
| 10 | Methods: Sample Size | (a) Specify the total number of animals used in each experiment, and the number of animals in each experimental group. | Methods |
| | | (b) Explain how the number of animals was arrived at. Provide details of any sample size calculation used. | Methods |
| | | (c) Indicate the number of independent replications of each experiment, if relevant. | Methods |
| 11 | Methods: Allocating Animals to Experimental Groups | (a) Give full details of how animals were allocated to experimental groups, including randomisation or matching if done. | Methods |
| | | (b) Describe the order in which the animals in the different experimental groups were treated and assessed. | |
| 12 | Methods: Experimental Outcomes | Clearly define the primary and secondary experimental outcomes assessed (e.g. cell death, molecular markers, behavioural changes). | Methods |
| 13 | Methods: Statistical Methods | (a) Provide details of the statistical methods used for each analysis. | Methods |
| | | (b) Specify the unit of analysis for each dataset (e.g. single animal, group of animals, single neuron). | Methods |
| | | (c) Describe any methods used to assess whether the data met the assumptions of the statistical approach. | Methods |

| | | | |
|---|---|---|---|
| 14 | Results: Baseline Data | For each experimental group, report relevant characteristics and health status of animals (e.g. weight, microbiological status, and drug or test naïve) prior to treatment or testing (this information can often be tabulated). | NA |
| 15 | Results: Numbers Analyzed | (a) Report the number of animals in each group included in each analysis. Report absolute numbers (e.g. 10/20, not 50%). | Results |
| | | (b) If any animals or data were not included in the analysis, explain why. | NA |
| 16 | Results: Outcomes and Estimation | Report the results for each analysis carried out, with a measure of precision (e.g. standard error or confidence interval). | Results |
| 17 | Results: Adverse Events | (a) Give details of all important adverse events in each experimental group. | NA |
| | | (b) Describe any modifications to the experimental protocols made to reduce adverse events. | |
| 18 | Discussion: Interpretation/ Scientific Implications | (a) Interpret the results, taking into account the study objectives and hypotheses, current theory and other relevant studies in the literature. | Discussion |
| | | (b) Comment on the study limitations including any potential sources of bias, any limitations of the animal model, and the imprecision associated with the results. | |
| | | (c). Describe any implications of your experimental methods or findings for the replacement, refinement or reduction (the 3Rs) of the use of animals in research. | |
| 19 | Discussion: Generalisability/ Translation | Comment on whether, and how, the findings of this study are likely to translate to other species or systems, including any relevance to human biology. | Discussion |
| 20 | Discussion: Funding | List all funding sources (including grant number) and the role of the funder(s) in the study. | Acknowledge-ments |

NA = not applicable.

**Fig. S2.** Isolation of Epithelial Cells from Porcine Pancreas

Protocol was adapted in part from techniques used to isolate pancreatic ductal epithelial cells (*Transplant Proc* 2009;41(4):1363-1366; PMID: 19460560). Total time from tissue explant to 10 cm dishes establishment should be in range of one month.

1. Isolate ducts from mass tissue removing as much of the acinar and islet cell tissue as possible in a sterile hood.
2. Mince up the tissue with a razor blade into 1mm$^3$ pieces and add 10 mg/mL of collagenase I plus 100 ug/mL DNase I.
3. Incubate cells at 37°C for 30 min and do mechanical dissociation with a 10 mL pipette every 10 min.
4. Centrifuge at 4°C (500 *g* x 5 min) to pellet cells and decant supernatant.
5. Resuspend in 10 mL of red blood cell lysis buffer ( and incubate for 10 min at room temp, then add 10 mL of media (DMEM + 5%FBS) and spin cells down again.
6. Take the cell pellet and resuspend with media and pass through a 70 μm nylon sieve (Corning™ Sterile Cell Strainers, Thermo Fisher Scientific, cat. no. 07-201-431).
7. Centrifuge cells again at 500 *g* x 5 min and decant supernatant.
8. Resuspend cells in 3 mL whole media and plate 1 mL in each well of a porcine gelatin coated 6 well plate.
9. After 4 hours take the media off the well and put in new well in 6 well plate to help remove fibroblast contamination.
10. Incubate cells overnight and the next day wash the wells with PBS and add fresh media (half DMEM, half ECM).
11. After 4 days epithelial cobblestone growth will start to appear and the epithelial cells will start to expand.
12. The well with the least amount of contaminating cells is used for further studies.
13. If fibroblasts are present the wells are trypsinized for 2-3 min and then washed with PBS x 2 to remove the fibroblasts. Remaining epithelial cells will expand from this and can then be plated in a 10 cm dish for expansion.
14. When a 10 cm dish is confluent the epithelial cells will be plated back into a 2- 6 well plates for immunocytochemistry and cellular transformation.

A

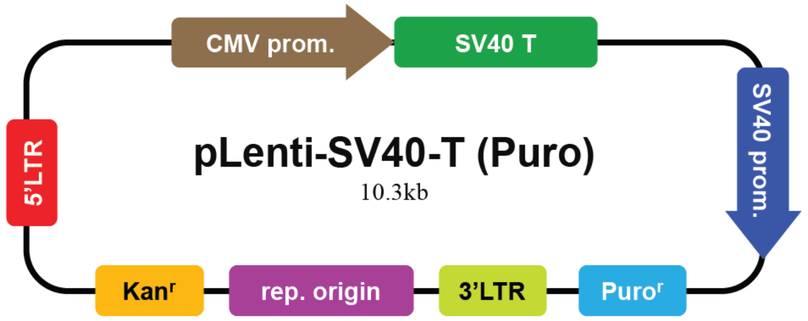


B

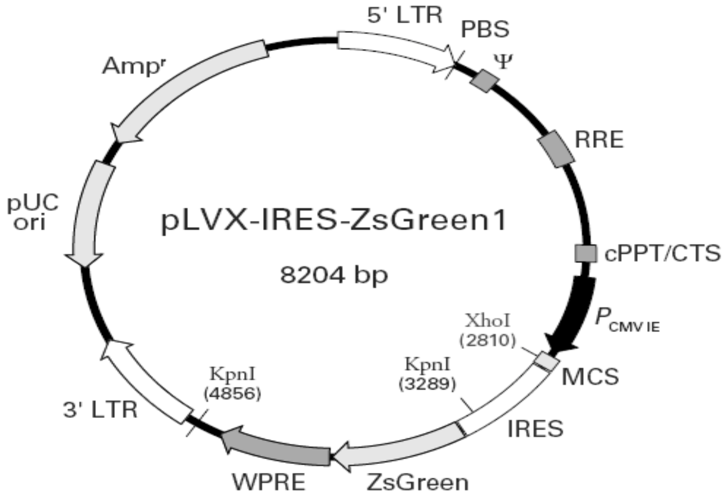


**Fig. S3**. Vector maps. (A) pLenti-SV40-T, from www.abmgood.com, cat. no. LV613. (B) pLVX-IRES-ZsGreen1, from www.takarabio.com, cat. no. 632187.

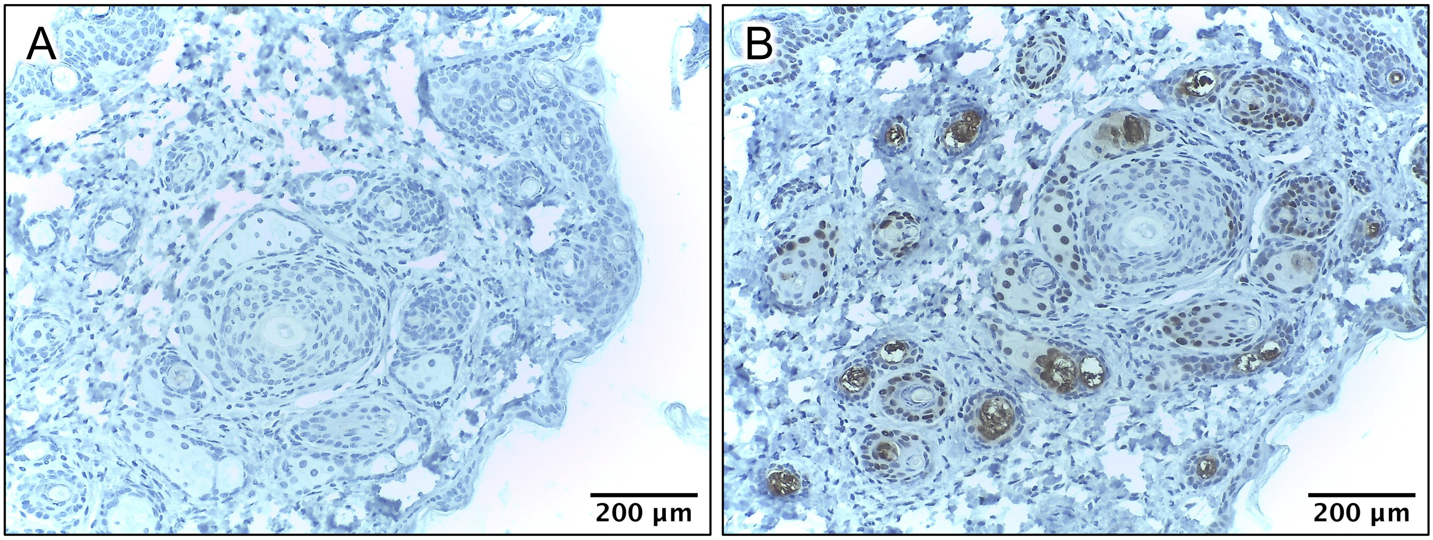


**Fig. S4**. Immunohistochemistry of p21 in nude mouse dermis/epidermis (p21 positive control). (A) Negative control (all reagents except no primary antibody). (B) p21.

**Table S1.** Responses to the NIH Principles and Guidelines for Reporting Preclinical Research.

| Principle/Guideline | Response | Location |
|---|---|---|
| **Standards**<br>Encourage the use of community-based standards (such as nomenclature standards and reporting standards like ARRIVE), where applicable. | ARRIVE | Fig. S1 |
| **Replicates**<br>Require that investigators report how often each experiment was performed and whether the results were substantiated by repetition under a range of conditions. Sufficient information about sample collection must be provided to distinguish between independent biological data points and technical replicates. | Done (benchtop assays) | Materials and Methods; Figure Legends |
| **Statistics**<br>Require that statistics be fully reported in the paper, including the statistical test used, exact value of N, definition of center, dispersion and precision measures (e.g., mean, median, SD, SEM, confidence intervals). | Done (benchtop assays & murine experiments) | Materials and Methods; Figure Legends |
| **Randomization**<br>Require authors to state whether the samples were randomized and specify method of randomization, at a minimum for all animal experiments. | Done (murine experiments) | Materials and Methods |
| **Blinding**<br>Require authors to state whether experimenters were blind to group assignment and outcome assessment, at a minimum for all animal experiments. | Done (pathologist evaluation of murine data) | Materials and Methods |
| **Sample-Size Estimation**<br>Require authors to state whether an appropriate sample size was computed when the study was being designed and include the | Done (murine experiments) | Materials and Methods |

| | | |
|---|---|---|
| statistical method of computation. If no power analysis was used, include how the sample size was determined. | | |
| **Inclusion and exclusion criteria**<br><br>Require authors to clearly state the criteria that were used for exclusion of any data or subjects. Include any similar experimental results that were omitted from the reporting for any reason, especially if the results do not support the main findings of the study. Describe any outcomes or conditions that were measured or used and are not reported in the results section. | NA | |
| **Data and Material Sharing**<br><br>Stipulate, at the minimum, that all datasets on which the conclusions of the paper rely must be made available upon request (where ethically appropriate) during consideration of the manuscript (by editors and reviewers) and upon reasonable request immediately upon publication.<br><br>Recommend deposition of datasets in public repositories, where available. Datasets in repositories should be bidirectionally linked to the published article in a way that ensures proper attribution of data production.<br><br>Encourage presentation of all other data values in machine readable format in the paper or its supplementary information. Require materials sharing after publication.<br><br>Encourage sharing of software and require at the minimum a statement in the manuscript describing if software is available and how it can be obtained. | Done | Acknowledgements |
| **Biological Variables**<br><br>Sex, age, weight, and underlying health conditions, are often critical factors affecting health or disease. In particular, sex is a biological | Done | Materials and Methods |

| | | |
|---|---|---|
| variable that is frequently ignored in animal study designs and analyses, leading to an incomplete understanding of potential sex-based differences in basic biological function, disease processes and treatment response.<br><br>Explain how relevant biological variables, such as the ones noted above, are factored into research designs, analyses, and reporting in vertebrate animal and human studies. Strong justification from the scientific literature, preliminary data or other relevant considerations must be provided for applications proposing to study only one sex. | | |
| **Authentication**<br><br>Key biological and/or chemical resources include, but are not limited to, cell lines, specialty chemicals, antibodies and other biologics.<br><br>Briefly describe methods to ensure the identity and validity of key biological and/or chemical resources used in the proposed studies. These resources may or may not be generated with NIH funds and:<br>• May differ from laboratory to laboratory or over time;<br>• May have qualities and/or qualifications that could influence the research data;<br>• Are integral to the proposed research. | Done | Materials and Methods |

Principles and Guidelines reproduced from online NIH resources for improving rigor, reproducibility, and transparency in biomedical research: (1) Principles and Guidelines for Reporting Preclinical Research. 17 December 2017. Available from: https://www.nih.gov/research-training/rigor-reproducibility/principles-guidelines-reporting-preclinical-research; (2) Enhancing Reproducibility through Rigor and Transparency. June 9, 2015. http://grants.nih.gov/grants/guide/notice-files/NOT-OD-15-103.html.

**Table S1.** DNA sequences used for mutagenesis, cloning.

| Gene | Purpose | Sequence |
|---|---|---|
| *KRAS*$^{G12D}$ | Clon | F: CTCGAGGCCGCCACCATGACTGAATATAAACTTGTGGTAGTTGGAGCTG<br>R: CTGCAGTTACATAATTATACACTTTGTC |
| *KRAS*$^{G12D}$ | PCR | F: CTTGTGGTAGTTGGAGCTG<br>R: CTCATGTACTGGTCCCTCATTG |

Mut = mutagenesis; Clon = cloning; F = forward sequence; R = reverse sequence

**Table S3.** Information on antibodies used for immunoblotting, immunofluorescence, and immunohistochemistry.

| Antigen | Product No. | Company | Website | Type | Host | Clone number |
|---|---|---|---|---|---|---|
| Cytokeratin 19 | ab7754 | abcam | Abcam.com | monoclonal | mouse | A53-B/A2 |
| Mutant $KRAS^{G12D}$ | GTX132407 | Genetex | Genetex.com | polyclonal | rabbit | |
| SV40T | ab16879 | abcam | Abcam.com | monoclonal | mouse | |
| Pan-Cytokeratin | ARG56128 | Arigo Biolaboratories | Arigobio.com | monoclonal | mouse | NA |
| Actin | AC026 | Abclonal | Abclonal.com | monoclonal | rabbit | |
| p21 | ab188224 | abcam | Abcam.com | monoclonal | rabbit | |
| PCNA | ab29 | abcam | Abcam.com | monoclonal | mouse | |

NA = not applicable.
